# Oxygen loss compromises the survival and cognition of a coastal cephalopod

**DOI:** 10.1101/2023.06.03.543560

**Authors:** Mélanie Court, Marta Macau, Maddalena Ranucci, Tiago Repolho, Vanessa Madeira Lopes, Rui Rosa, José Ricardo Paula

## Abstract

The ocean is undergoing deoxygenation and the spread of hypoxic areas. Ocean deoxygenation and standing levels of hypoxia are shrinking fundamental niches, particularly in coastal areas, yet documented repercussions on species development and behavior are limited. Here, we tackled the impacts of deoxygenation (7 mg O2 L^-1^), mild hypoxia (nocturnal 5 mg O2 L^-1^), and severe hypoxia (2 mg O2 L^-1^) on cuttlefish (*Sepia officinalis*) development (hatching success, development time, mantle length) and behavior, i.e., ability to learn (associative-and socially), to camouflage, and to explore its surroundings spatially. We found that hypoxia yielded lower survival rates, smaller body sizes and inhibited predatory (increased latency to attack the prey) and anti-predator (camouflage) behaviors. Acute and chronic exposure to low oxygen produced similar effects on cognition (inability to socially learn, increased open-field activity levels, no changes in thigmotaxis). It is thus expected that, although cuttlefish can withstand oxygen limitation to a certain degree, expanding hypoxic zones will diminish current habitat suitability.

## Introduction

Anthropogenic greenhouse gas emissions and nutrient inputs from coastal activities are precipitating increasingly severe changes in ocean’s oxygenation, a phenomenon often dissimulated by high oxygen temporal and spatial variability (Breitburg et al., 2018; Schmidtko et al., 2017). The ocean has been evincing a continuous decline in dissolved oxygen of 0.5-3.3% per decade since 1970 (Cooley et al., 2022). This trend is expected to provoke a reduction in the ocean’s oxygen content of 4.1% to 11.2% by 2100, according to the Shared Socioeconomic Pathways 2-4.5 and 5-8.5, respectively (Cooley et al., 2022; Kwiatkowski et al., 2020). On the one hand, ocean oxygen loss emerges from the warming of the ocean, which reduces oxygen solubility in water and boosts biological O2; on the other hand, an increase in stratification and changes in circulation diminish ocean ventilation (Schmidtko et al., 2017).

In addition, oxygen-minimum zones (OMZ) are expanding, shoaling the depth of hypoxic waters (Breitburg et al., 2018; Keeling et al., 2010). Climate shifts further accentuate periodic hypoxia (assumed at 2 mg O_2_ L^-1^), often leading to behavioral changes and mortality in nekton-benthic communities (Diaz & Rosenberg, 1995). Furthermore, coastal areas and enclosed bodies of water are particularly susceptible to eutrophication-induced hypoxia caused by excess nutrient input from anthropogenic activities (Cai et al., 2011; Melzner et al., 2012). Hypoxia is likely to heighten during the night due to the absence of photosynthesis (Summers & Engle, 1993; Newton et al., 2010). Oxygen limitation affects aerobic processes, such as energy metabolism (Rosa and Seibel, 2008, 2010). Larger-bodied organisms have higher metabolic requirements and sustain a higher oxygen demand (Deutsch et al., 2020). Lower oxygen concentrations are often encountered in warmer areas, where organisms face higher metabolic requirements. It is often the driving factor behind distributional population shifts (Deutsch et al., 2020).

Despite being highly predated upon, coleoid cephalopods, i.e., octopuses, cuttlefish, and squid, are highly active predators (Amodio et al., 2019) and require significant oxygen provision (Wells, 1990). Oxygen loss can thus cause depletion of energy and amino acid reserves (Jiang et al., 2020) and is limiting to the performance and survival of cephalopods (O’Dor & Webber, 2011; Pörtner, 2002; Seibel, 2016).

Cephalopods exhibit advanced encephalization, possessing the highest brain-body mass ratios among invertebrates (Hermann, 2017). This may confer them behavioral plasticity or adaptability to changing environments (Borrelli, 2007; Pollen et al., 2007; Vitti, 2013), although this is challenging to confirm in this group (Ponte et al., 2021). Cephalopod vertebrate-like cognitive abilities provide them with a vast learning capability (Hanlon & Messenger, 2018c). Regarding cuttlefishes, although no learning was registered regarding hunting techniques (Boal et al., 2000) nor danger avoidance (Huang & Chiao, 2012) in adult *Sepia officinalis*, newborns do learn from conspecifics (Sampaio et al., 2020). This behavior is an advantage of social groupings, as it saves time and energy by preventing trial-and-error learning (Coussi-Korbel & Fragaszy, 1995). The opportunity for social learning arises during spawning, which indicates that newborns also exhibit gregariousness (Drerup & Cooke, 2021; Hanlon & Messenger, 2018b). Further, juveniles have been found constituting shoals during migration (Drerup & Cooke, 2021), suggesting that cuttlefish developed a general type of social cognition, which can be useful in both social and non-social environments (Varela et al., 2020).

Cuttlefish rely on camouflage for protection in the juvenile stage and intra-and interspecific communication in adulthood (Hanlon & Messenger, 2018a). Their highly precise visual and neurological systems allow them to be particularly effective predators (Budelmann, 1996; Villanueva et al., 2017) and lend them a unique ability to instantly change body patterns and, thus, camouflage themselves from predators (Hanlon & Messenger, 2018a). To achieve this, cuttlefish either emulate background texture and colors (uniform or mottle patterns) or produce disruptive patterns, which generate false edges and conceal the animal’s true body outline (Hanlon & Messenger, 2018a). Further, the visual system is easily impaired by oxygen deficiency (McCormick et al., 2019). Due to their reliance on vision for camouflage, cuttlefish are especially prone to impacts driven by oxygen limitation.

Despite being the least studied, hypoxic events produce steeper repercussions on marine organisms than ocean warming and acidification combined (Sampaio et al., 2021). This highlights the need for a comprehensive audit of the consequences of these events on marine biota (Vaquer-Sunyer & Duarte, 2006; Cooley et al., 2022; Sampaio et al., 2021), as well as the impacts of continuous oxygen limitation (Borges et al., 2022). Further, direct effects on survival and growth should be extensively understood. However, sub-lethal effects, such as behavioral and cognitive deleterious responses, can also have long-lasting impacts on populations. Here, we analyze the developmental and behavioral responses to oxygen limitation in the commercially-important European cuttlefish S. officinalis. More precisely, we investigated survival, body size, asocial and social learning capacity, anti-predator behavior (camouflage contrast), and foraging-inhibiting responses (activity levels, thigmotaxis) of cuttlefish hatchlings to chronic (deoxygenation, 7 mg O2 L^-1^) and acute exposure to oxygen limitation - mild hypoxia (nocturnal 5 mg O_2_ L-1 [least conservative hypoxia estimate; Vaquer-Sunyer & Duarte, 2008]) and severe hypoxia (nocturnal 2 mg O2 L-1). These findings will provide valuable insights into the capacity of an ecologically and economically important species to withstand a largely unexplored but increasingly dangerous stressor.

## 2. Methods

### 2.1. Cuttlefish rearing

Early-stage *S. officinalis* eggs were collected on the Tagus estuary (Lisbon Area, Portugal) by local fishermen and transported to Laboratório Marítimo da Guia (LMG, Cascais). After laboratory acclimation (three days), the eggs were placed in twelve 9-L plastic tanks. The eggs (429) were distributed randomly between four treatments, with a higher sample size in Normoxia to account for additional cuttlefish required for learning trials:

- Normoxia: 8 mg O_2_ L^-1^ (153 eggs, 51 per tank)
- Deoxygenation: 7 mg O_2_ L^-1^ (99 eggs, 33 per tank)
- Mild hypoxia (MH): nocturnal 5 mg O_2_ L^-1^ (93 eggs, 31 per tank)
- Severe hypoxia (SH): nocturnal 2 mg O_2_ L^-1^ (84 eggs, 28 per tank)

Natural seawater (NSW) was pumped directly from the sea, filtered through a 1- micrometre mesh filter, and sterilized by a 12-W UV-sterilizer (Vecton 120 Nano, TM-Iberia, Lisbon, Portugal). Circulating NSW was continuously renewed via a drip system and oxygenated through air stones connected to an air compressor (Medo Blower LA- 120A, Nitto Kohki, Japan). NSW in each tank was renewed every 30 minutes (water flow of approx. 250 mL min^-1^). The tanks were illuminated by LED 8-W lights, under a 14 h light / 10 h dark photoperiod. Temperature was maintained with temperature controllers (HX-W3002, accuracy ± 0.1°C, hysteresis 0.3°C) connected to 150-W water heaters (Eheim GmbH & Co KG, Deizisau, Germany) and a water chiller (TK500, Teco, Ravenna, Italy).

Oxygen levels were regulated independently in each system. Oxygen concentration was controlled by solenoid valves connected to an Arduino Mega controller (Mucha, 2023; hysteresis set at 0.2 mg O_2_ L^-1^), which read oxygen levels through an optometer (PyroScience FireStingO2, accuracy ± 0.1 mg O_2_ L^-1^). Solenoid valves injected certified N_2_ gas (Air liquide, Algés, Portugal) into three cylindrical reservoirs to stabilize O_2_ levels (including air stones to recover oxygen levels when needed). Reservoirs fed oxygen-limited water to the tanks via a water pump (35 W; TMC, V2 Power Pump, 2150 L h^-1^). The water flow was adjusted manually daily through an acrylic flow meter (1-10 L min^-1^ range). Nitrogen injection in MH and SH treatments started at 00h00 and halted at 7h30 daily, regulated by a Profilux controlling system (3N GHL, Kaiserslautern, Germany), whilst N_2_ injection in the Deoxygenation treatment was continuous. MH and SH treatments required 80 minutes and 120 minutes, respectively, to recover to 100% oxygen saturation after N_2_ injection cessation. An air pump (Aqua One, Stellar 380D Series II) was used to hasten O_2_ recovery.

Temperature and oxygen levels (oximeter VWR DO220, accuracy ± 0.1 mg O_2_ L^-1^, ± 0.3°C), salinity (Hanna refractometer, accuracy ± 1 ppt), and pH (pH meter VWR pHenomenal, accuracy ± 0.005), were monitored twice daily in each tank (at 7h30 during the nocturnal treatment [Supp. Table 1] and 10h00, after oxygen recovery [Supp. Table 2]). Total ammonia nitrogen, nitrite, and nitrate levels were monitored weekly using colorimetric tests (TropicMarin, Hünenberg, Switzerland) and kept below detectable levels.

Hatchlings were fed laboratory-reared sub-adult amphipods *Gammarus locusta* three days post-hatching, and the day before the social learning trial. Upon hatching, cuttlefish were placed in individual cups (with four netted exits for water circulation), labeled with the hatching date and the individual ID, and kept in their tank. Hatching success was considered as the number of hatchlings over the number of fertilized eggs. Mantle length was extracted through ImageJ using photographs from camouflage trials (using arena size as a reference). Mantle lengths were calculated by one observer (MM), and randomly chosen photographs (20%) were analyzed by a second observer (MC).

### 2.2. Asocial and social learning

To examine their ability to associatively learn through instrumental (asocial learning with response-reinforcer – lack of reward, Cartron et al., 2013) and observational (social learning with response-reinforcer) learning (Heyes, 1994), cuttlefish underwent prawn-in-a-tube trials, with 140 demonstrators and 140 observers. From three to seven days post-hatching, they were placed in an experimental arena (13.5 x 5.5 x 4 cm acrylic rectangular container and a see-through partition in the middle). Each arena contained a demonstrator cuttlefish (exposed to the amphipod *G. locusta* in an inaccessible cylindrical glass tube) and an observer, able to observe the unsuccessful attack attempts of the demonstrator through the transparent divider (**Figure 1**). All outward-facing walls were previously covered with black plastic to prevent cuttlefish from seeing neighboring individuals.

**Figure 1.**
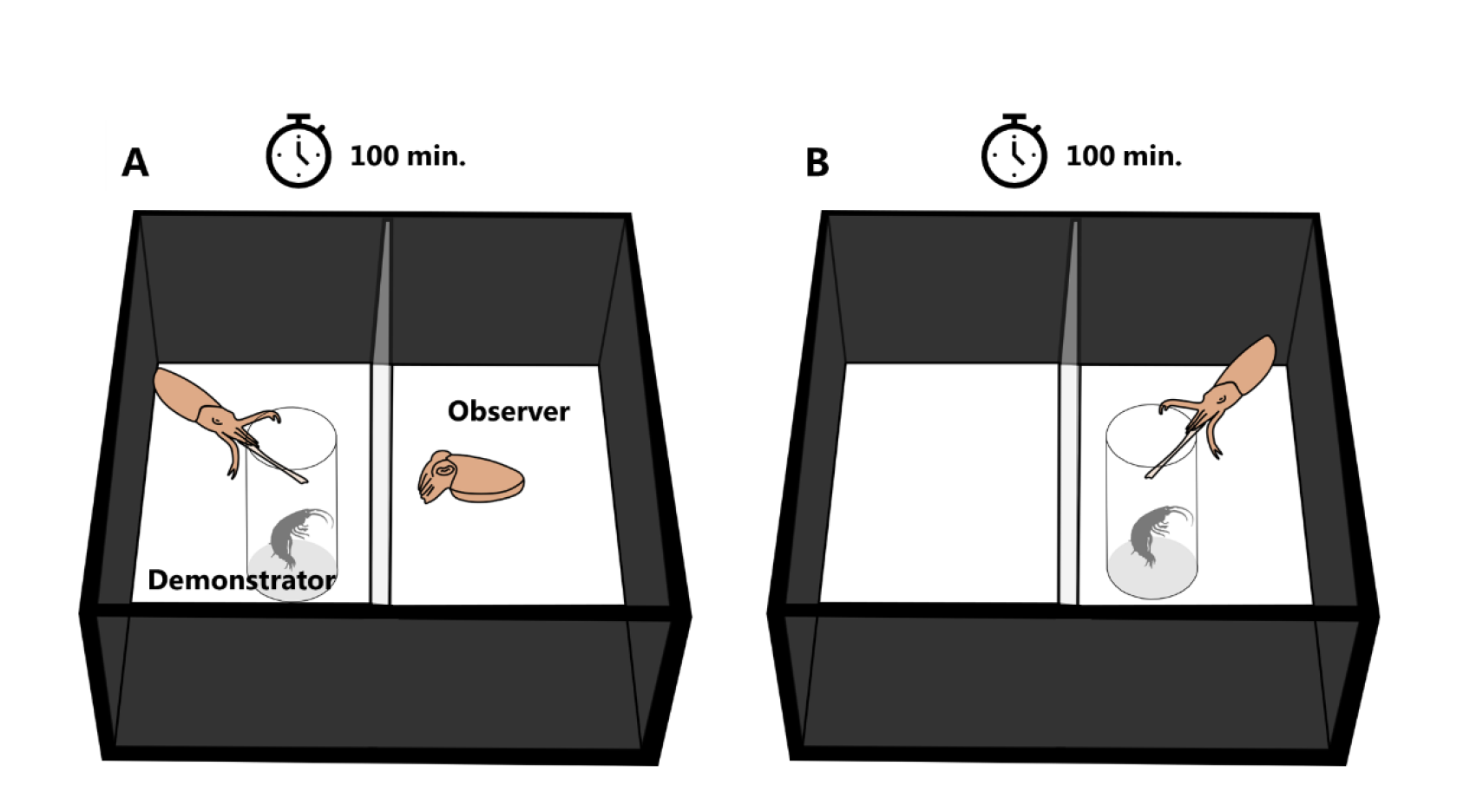
Experimental setup to test associative and social learning. A) The amphipod-in-a-tube is placed in the demonstrator compartment for a hundred minutes, at 10- minute intervals. B) After thirty minutes, the amphipod-in-a-tube is placed in the observer compartment for another 100 minutes, at 10-minute intervals

The arena was filled with approx. 200 mL of water from the cuttlefish’s tank prior to each trial. Video cameras (Sony Handycam DCR-SR78 and Canon Legria HF R56) filmed from a 90° angle, covering several arenas. After being placed in the arena with a black spoon, the two cuttlefish underwent a sixty-minute acclimation period. Filming started upon exposure of the demonstrator cuttlefish to the prawn-in-the-tube. Both cuttlefish were left to observe the prey for ten minutes, after which the tube was withdrawn. After an additional ten minutes, the amphipod was again placed in the demonstrator section. This procedure was repeated ten times. Following a thirty-minute break, the same steps were undertaken relative to the observer, i.e., with the prey tube in the observer section. To control for the effect of the treatments on demonstration, observers were paired with demonstrators from their own treatment and from normoxic conditions.

From video recordings, the latency to the first attack (up to 600 seconds) was registered according to the function (demonstrator or observer) and the treatment. The criterion for successful learning was defined as three consecutive trials wherein cuttlefish did not attack the prawn (Sampaio et al., 2020). Successful learning from the demonstrators was considered asocial learning, whereas learning from the observers was considered social learning. In addition, learning was evaluated through the latency to the first attack.

### 2.3. Exploratory-avoidance

To determine whether cuttlefish spatial exploration is inhibited by limited oxygen availability, an open-field test was performed following a protocol adapted from Hamilton et al. (2014). Two days after the learning trials, cuttlefish (N = 37) were transferred to a white circular arena filled with the cuttlefish’s treatment water (400 mL). A purple bottle cap was placed in the center to facilitate exploratory behavior, surrounded by an elevated platform. The arena was enveloped by white styrofoam to diffuse the light and separated from the observer via black curtains. The cuttlefish were recorded (LEGRIA HF R56 35 Mbps, Canon, Porto Salvo, Portugal) from above for twenty minutes, from the moment they were placed in the arena. Animal tracking data were extracted from the video recordings through the software ToxTrac v2.61 (Rodriguez et al., 2018) using the ToxId algorithm (Rodriguez et al., 2017).

### 2.4. Camouflage

To evaluate the chromatic component of camouflage, cuttlefish body pattern matching was compared to the substrate they were placed upon. The day after the learning trials, cuttlefish (N = 90) were placed in a white round arena using a black spoon, with the bottom covered with a 60% black and 40% white gravel mixture (Fishnet, Lisbon, Portugal) to induce the disruptive body pattern, or in the sand to induce the mottle pattern. The arenas were filled with approx. 400 mL of water from the cuttlefish tank. The order of presentation of the substrate type was alternated between trials. A random number displayed in the photographs was attributed to all cuttlefish to prevent observer bias in data analysis (blinding).

A video camera (GoPro Hero 3+, San Mateo, CA, USA) recorded the arena from a 75° angle for ten minutes following acclimation (until cuttlefish spent five seconds stationary) to register burying attempts and account for the locomotor component of camouflage (Hanlon & Messenger, 2018a). Photographs were taken remotely (Canon PowerShot G7X Mark II, white balance-calibrated, shutter speed 1/15, F-stop f/11, ISO 250, 1080p, 60 fps) at a 90° angle upon every camouflage pattern change or pattern intensification. Concurrently, the time of photograph shooting was registered using a chronometer.

Camouflage latency was considered as the time past acclimation until the photograph in which cuttlefish camouflaged in each substrate (strong mottle in sand and a dark uniform or disruptive pattern in gravel) was taken. The strength of disruptive patterns in gravel was evaluated through the difference between maximum and minimum pixel values (grayscale) in the frontal body plane (Court et al., (2022) adapted from Chiao et al., (2009)) read in the ImageJ 1.46r software (National Institute of Health, USA). Cuttlefish can be equally as well camouflaged when donning a dark uniform pattern instead of the classical disruptive. To account for this, minimum intensity values were used to assess whether cuttlefish resorted to dark uniform patterns. In sand, the mean and standard deviation of pixel intensities were used to evaluate the strength of the mottled pattern. A second researcher analyzed a subset of photographs (20%) to account for observer bias.

### 2.5. Statistical analyses

Statistical analyses were performed in the R software, PBC, version 4.1.2. Data exploration was performed following Zuur & Ieno (2016), using the HighstatLibV10 R library from Highland Statistics (Zuur et al., 2009). The level of significance was set at p < 0.05. Differences between measurements of camouflage intensities were checked through Pearson correlation tests (“corr.test”, package “corrplot”). All correlations were significant (p < 0.001, Table S7), with coefficients superior to 0.94.

#### 2.5.1. Hatching success and mantle length

Hatching over time was evaluated through time-to-event analyses, which right-censor data to account for true survival times being equal to or greater than those observed at the end of the study (Schober & Vetter, 2018). Since the incorporation of mixed effects did not permit testing the model’s assumptions, a Cox Proportional Hazards (CPH) regression model (“coxph”, R package “survival”) was fitted to hatching success data. Days to hatching and successful hatching (binary factor, 0/1) were considered as covariates. The model was fitted by maximum likelihood with treatment as the predictor variable (factor of four levels - N, DO, MH, and SH). The assumptions of the “coxph” model (proportional hazards, no over-influential observations and linearity of co-variates) were not met (Schoenfeld test, represented by a smoothed spline plot of the scaled Schoenfeld residuals against time), thus a “survdiff” model was fitted. Post-hoc multiple comparisons were performed (“pairwise_survdiff” function), and p-values were adjusted through Bonferroni-Hochberg corrections to avoid type I errors. The survival curve was plotted as a Kaplan-Meier plot (function “ggsurvplot” from the R package “survminer”).

To determine the influence of treatments on cuttlefish mantle length, a Linear Mixed Model (LMM) (identity link function) was fitted, using a Template Model Builder (function “glmmTMB” from the package “glmmTMB”; Brooks et al., 2017). Treatment was considered a four-level factor (fixed effect), with cuttlefish age and replicates (three-level factor) as random effects. A type II Wald chi-squared test (“Anova” function in the “car” package; Fox & Weisberg, 2018) was performed to assess the influence of treatments on mantle length. Post-hoc multiple comparisons (“emmeans” function in the ”emmeans” package) were performed, and p-values were adjusted through Tukey corrections. LMM assumptions (normality, homoscedasticity, and independence of residuals) were checked by plotting the model’s residuals.

#### 2.5.2. Learning

The difference in the proportion of successful learning between observers and demonstrators was evaluated through GLMM from the binomial family (logit link function) with function (demonstrator/observer) as the predictor variable, successful learnings (0/1) as the response, and the replicate as a random effect. The difference in learning rate was assessed through a poisson family GLMM, with the trial-at-learning as a predictor variable.

The effect of the demonstrator treatment on observer learning was inferred from a CPH regression model (“coxph” function). The model assumed the treatment of the demonstrator as a two-level factor (Normoxia or same treatment as observer) and trial-to-learn and learning success (0/1) as response variables. The proportional hazards assumption was met. Similarly, the effects of treatments on asocial and social learning (learning from the demonstrator and the observer, respectively) were evaluated through the “coxph” function, with treatment as a four-level factor.

A LMM (identity link function) was fitted to latencies to attack. Treatment and function were considered as fixed effects, and the replicate as random effect. The models best fitted to the data were selected through the AIC and included the treatment and trial number interaction.

#### 2.5.3. Exploratory-avoidance and camouflage performance

LMMs were fitted to the average speed and proximity to walls (time of permanence near the walls), with the replicate as a random effect. LMMs were fitted to camouflage latencies in gravel and in sand, as well as to pixel intensities in sand and gravel. Replicate and first substrate presented to the cuttlefish were considered random effects. GLMMs from the binomial family were fitted to burial attempts in the sand.

### 2.6. Ethical statement

All experiments were approved by the FCUL Animal Welfare Committee (ORBEA FCUL; permit 2/2022) as per National (DL 113/2013 and DL 1/2019) and EU legislation (Directive 2010/63/EU).

## 3. Results

### 3.1. Time-to-hatch and body size

Severe hypoxia (SH)-treated cuttlefish displayed markedly reduced hatching success (74.2%) relative to Normoxia (84.0%) and hatched significantly later (average 50.5 ± 12.3 days; Normoxia, 41.8 ± 8.9) (CPH, df = 270, p < 0.001; **Figure 2A**). In addition, SH-exposed cuttlefish hatched smaller than in Normoxia (LMM, t = 3.34, df = 92, p = 0.001; **Figure 2B**).

**Figure 2.**
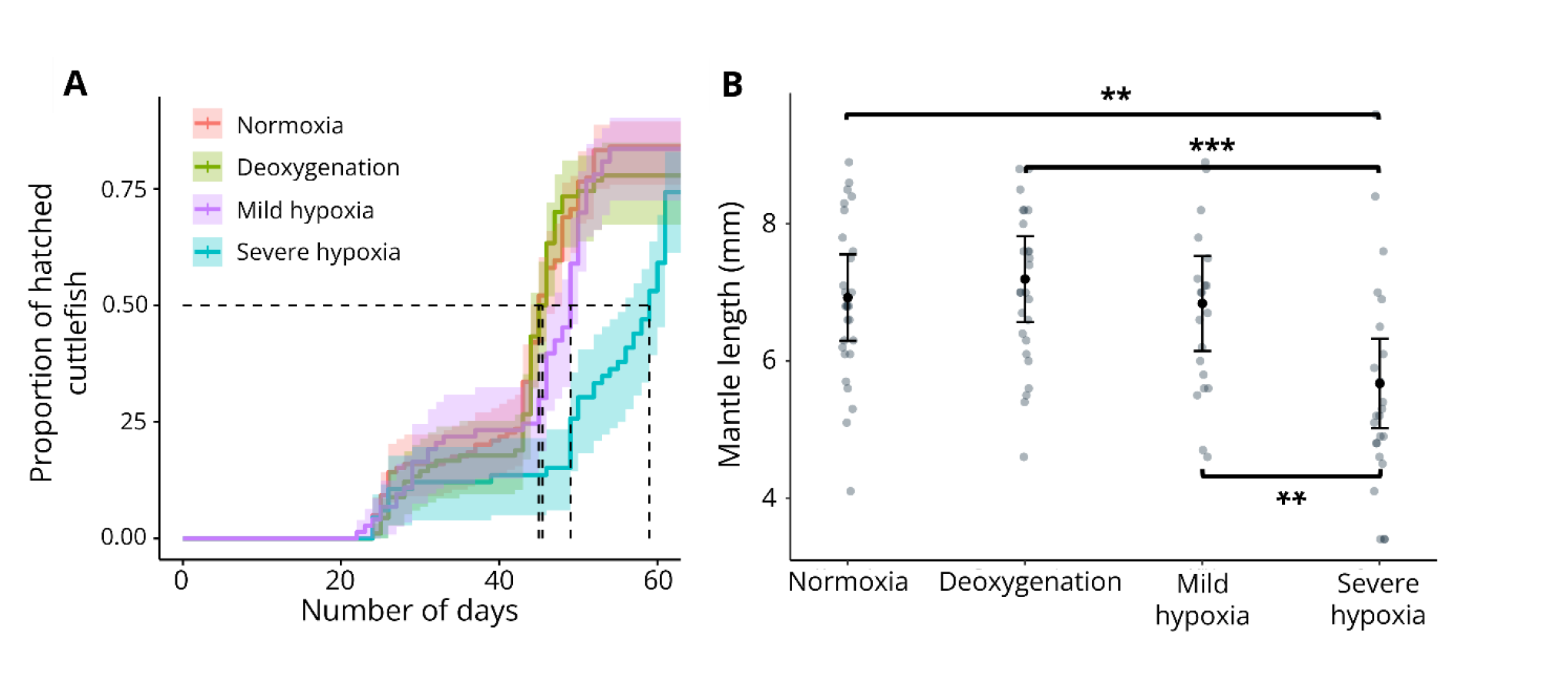
Indicators of developmental performance under the treatments Normoxia (8 mg O_2_ L^-1^), Deoxygenation (7 mg O_2_ L^-1^), Mild hypoxia (nocturnal 5 mg O_2_ L^-1^), and Severe hypoxia (nocturnal 2 mg O_2_ L^-1^). A) Cumulative proportion of hatched cuttlefish from egg capture date. B) Mantle length measured in three-to seven-day old cuttlefish. Lines in A) represent predicted proportions (Cox regression), and shaded areas depict 95% confidence intervals. Results in B) are expressed as estimated marginal means ± CI (95%). Dots in B) represent individual observations. Comparison significance levels: * p < 0.05, ** p < 0.01, *** p < 0.001

### 3.2. Asocial and social learning

Most cuttlefish fulfilled the criterion for learning. Observers learned in higher proportions (87.5%; **Figure 3B**) than demonstrators (61.4% of demonstrators, **Figure 3A**; superposed probabilities in **Figure S1**) (Wald chi-squared, χ^2^ = 22.5, df = 1, p < 0.001; statistical outputs shown in **Tables S3** and **S4**). Moreover, observers learned on average faster (average trial 3.4 ± 3.2) than demonstrators (trial 7.9 ± 2.3) (Wald chi-squared, χ^2^ = 194.0, df = 1, p < 0.001). The treatment of the demonstrator had no effect on learning from the observer (Wald chi-squared, χ^2^ = 0.0, df = 1, p = 0.9; **Figure S2**).

**Figure 3.**
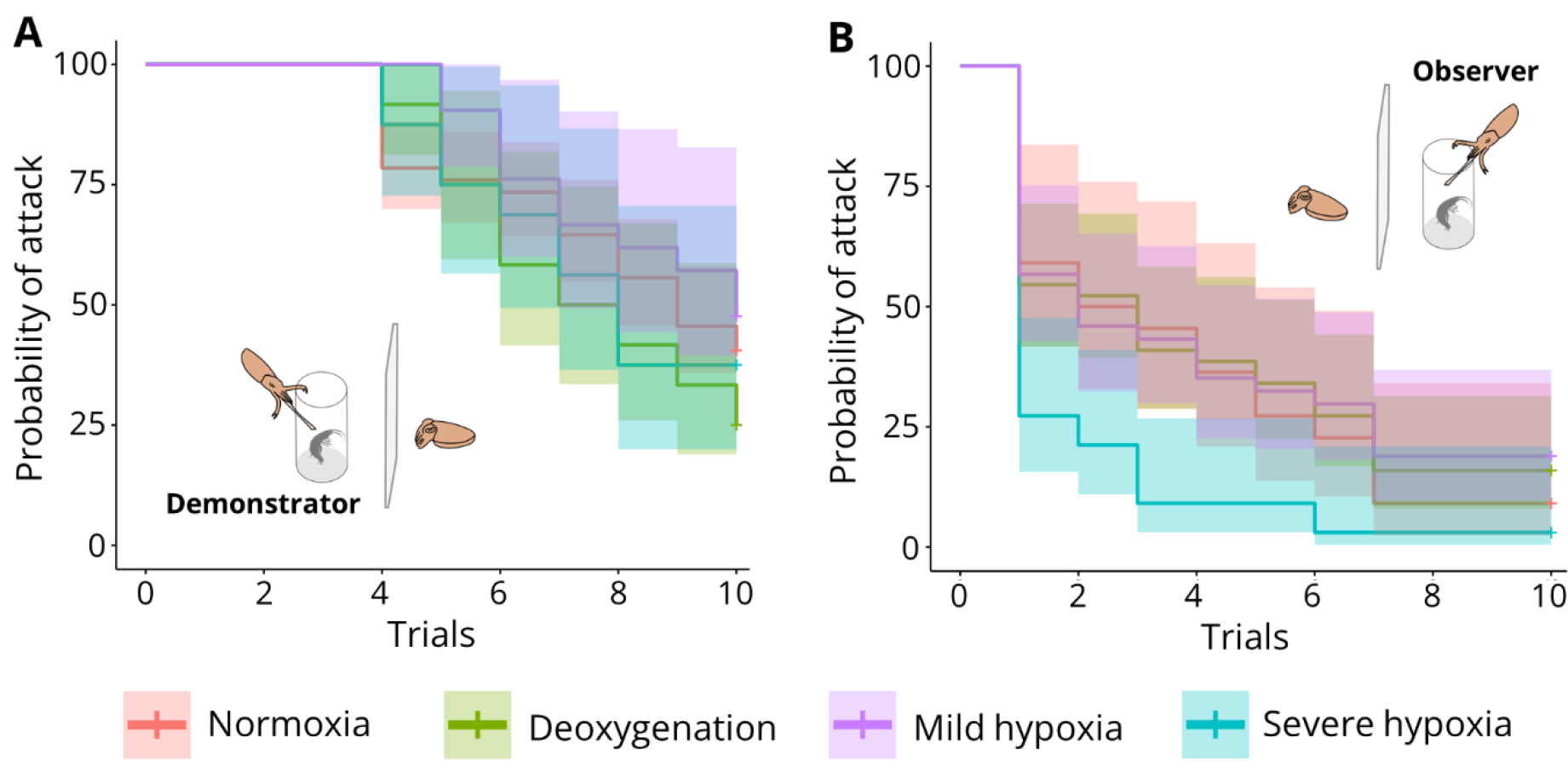
Probability of attacking the inaccessible amphipod across trials, obtained through time-to-event analysis, under the treatments Normoxia (8 mg O_2_ L^-1^), Deoxygenation (7 mg O_2_ L^-1^), Mild hypoxia (nocturnal 5 mg O_2_ L^-1^) and Severe hypoxia (nocturnal 2 mg O_2_ L^-1^). Lines represent predicted proportions (Cox regression), and shaded areas depict 95% confidence intervals

The treatments did not affect asocial learning (Wald chi-squared, χ^2^ = 2.9, df = 3, p = 0.4). Observers had 31.8% less probability of attacking under SH compared to Normoxia (CPH, z = 2.3, df = 135, p = 0.021). However, only 27.3% of these cuttlefish ever attacked (**Table S5**). Moreover, increased latency to the first attack from observers was observed solely in control (LMM, t = 50.8, df = 267, p = 0.022; **Figure 4**). Considering learning as three consecutive trials without attacking after failing at least once, demonstrator treatments had no effect on learning from observers (Wald chi-squared, χ^2^ = 2.9, df = 3, p = 0.4). The lack of attack attempts was reflected in an increased latency to attack in SH (LMM, t = - 3.5, df = 273, p = 0.003).

**Figure 4.**
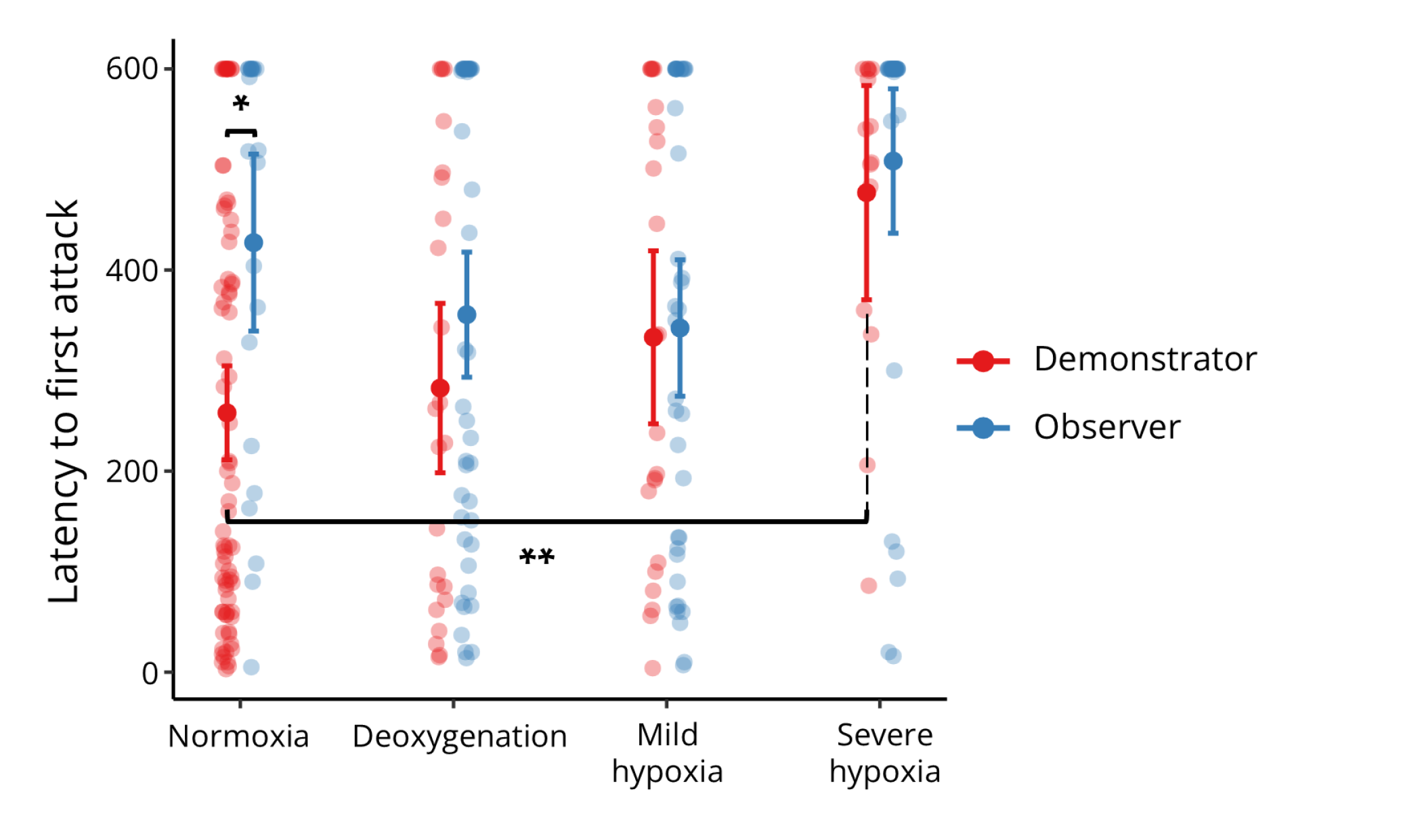
Latency to the first attack of the inaccessible amphipod in demonstrator (red) and observer (blue) cuttlefish under the treatments Normoxia (8 mg O_2_ L^-1^), Deoxygenation (7 mg O_2_ L^-1^), Mild hypoxia (nocturnal 5 mg O_2_ L^-1^) and Severe hypoxia (nocturnal 2 mg O_2_ L^-1^). Results are expressed as estimated marginal means ± CI (95%). Dots represent individual observations. Comparison significance levels: * p < 0.05, ** p < 0.01, *** p < 0.001

### 3.4. Exploratory-avoidance

To evaluate exploratory-avoidance, speed (**Figure 5A**), spatial exploration of the arena (**Figure S3**), and closeness to its walls (thigmotaxis; **Figure 5B**) were quantified (statistical outputs in **Table S6**). Deoxygenation-and Mild hypoxia (MH)-treated cuttlefish displayed increased average speeds than Normoxic cuttlefish (LMM, t = −3.29, p = 0.013; t = −4.303, p = 0.001). This was reflected in higher spatial exploration (LMM, t = −3.16, df = 31, p = 0.018; t = −3.57, p = 0.006; respectively). Oxygen limitation did not affect the proportion of time spent near the walls (Wald chi-squared, χ^2^ = 5.5, df = 3, p = 0.060).

**Figure 5.**
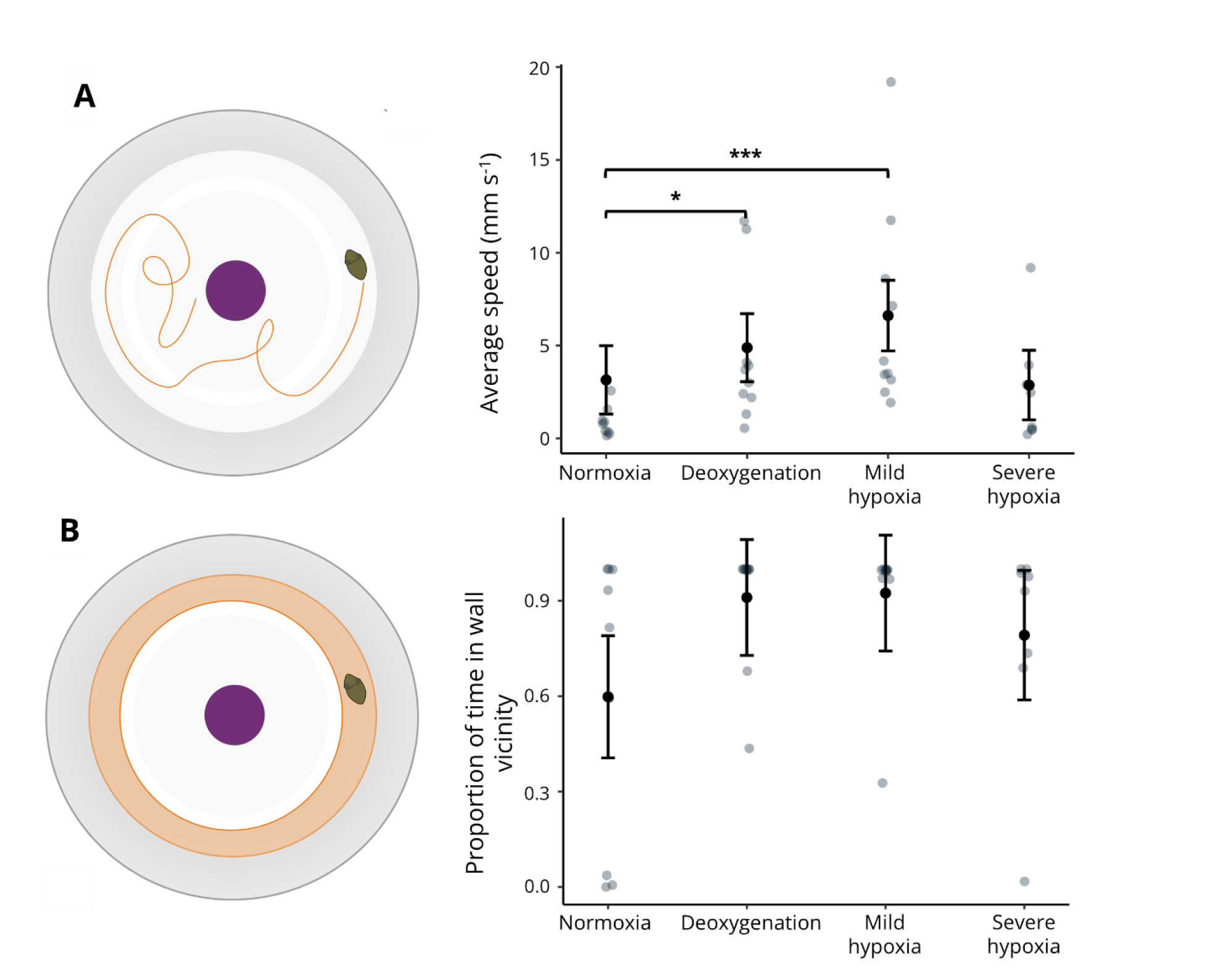
Parameters extracted from an open-field test relative to the treatments Normoxia (8 mg O_2_ L^-1^), Deoxygenation (7 mg O_2_ L^-1^), Mild hypoxia (nocturnal 5 mg O_2_ L^- 1^), and Severe hypoxia (nocturnal 2 mg O_2_ L^-1^). A) Average speed. B) Percentage of time spent in wall vicinity (highlighted in orange). Results expressed in estimated marginal means ± CI (95%). Dots represent individual observations. Comparison significance levels: * p < 0.05, ** p < 0.01, *** p < 0.001

### 3.5. Camouflage

The range of pixel intensities along the cuttlefish’s frontal body plane in gravel, indicating the strength of the disruptive pattern, was lower in SH-treated cuttlefish relative to Normoxia (LMM, t = 2.70, df = 85, p = 0.041) (**Figure 6A**; statistical outputs shown in **Tables S7 and S8**). The minimum pixel intensity, i.e., the ability to produce dark tones, was higher in cuttlefish from SH compared to control (LMM, t = −3.90, df = 85, p = 0.001; **Figure 6B**). The pixel range of cuttlefish in sand was not affected by the treatments (Wald chi-squared, χ^2^ = 0.39, df = 3, p = 0.941; **Figure 6C**), nor was the minimum pixel intensity (χ^2^ = 0.20, df = 3, p = 0.978; **Figure 6D**).

**Figure 6.**
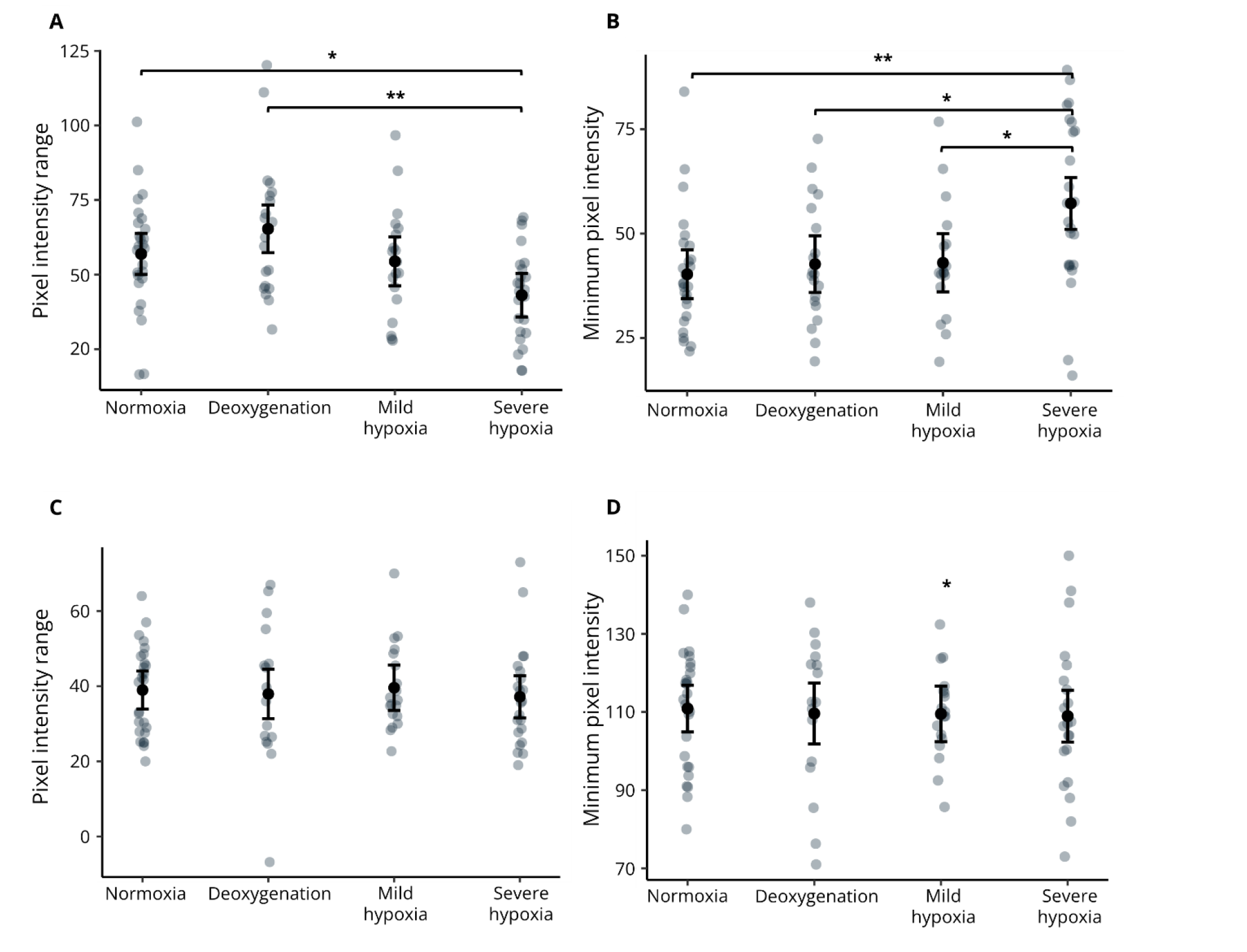
Camouflage performance relative to the treatments Normoxia (8 mg O_2_ L^-1^), Deoxygenation (7 mg O_2_ L^-1^), Mild hypoxia (nocturnal 5 mg O_2_ L^-1^), and Severe hypoxia (nocturnal 2 mg O_2_ L^-1^). A) Difference between maximum and minimum pixel intensities in the frontal body plane of the cuttlefish in gravel. B) Minimum pixel intensities along the frontal body plane in gravel. C) Difference between maximum and minimum pixel intensities in the frontal body plane of the cuttlefish in sand. D) Minimum pixel intensities along the frontal body plane in sand. Results are expressed in estimated marginal means ± CI (95%). Dots represent individual observations. Comparison significance levels: * p < 0.05, ** p < 0.01, *** p < 0.001

Latencies to camouflage were not affected by the treatments (**Figure S4**) in sand (Wald chi-squared, χ^2^ = 9.6, df = 3, p = 0.31) nor in gravel (χ^2^ = 2.6, df = 3, p = 0.053). Cuttlefish exposed to MH oxygen levels attempted to bury themselves in sand more frequently than those exposed to Deoxygenation (GLMM, t = −2.73, df = 93, p = 0.038).

## 4. Discussion

Here we investigated the effects of early-life exposure to acute and chronic oxygen limitation on the development and cognition of *Sepia officinalis*. We found that exposure to Severe Hypoxia (2 mg O_2_ L^-1^) induced fewer hatchings and delayed development. The mass-specific metabolic rates of developing organisms are higher than those of adults, which exposes them to higher oxygen-deficiency risk (Sibly et al., 2015; Somero et al., 2016). In fact, on average, early life stages of marine species survive 64% less to hypoxia than adult individuals. In particular, cuttlefish embryos encounter difficulties in gas exchange as they are enveloped in a gelatinous capsule (Rosa et al. 2013). Thus, the subsequent increase in metabolic and energetic requirements under Severe hypoxia likely delayed development, stunted growth and reduced survival.

### Inhibitory control and motivation to hunt

The prawn-in-a-tube test has been extensively used to assess the cognitive abilities of cephalopods and the neurological foundation of learning (Cartron et al., 2013). In the said test, cuttlefish learn to inhibit predation (aversive learning) given the absence of reward. Thus, the suppression of predatory behavior could be attributed to habituation to the stimulus, wherein response evocation wanes over the number of trials in the absence of stimulus reinforcement (Messenger, 1973). This is unlikely to occur in the cuttlefish as dishabituation was not found after learning in the prawn-in-a-tube test (Agin et al., 2006; Bowers et al., 2020; Messenger, 1973; Purdy et al., 2006). Observer cuttlefish achieved greater learning success and faster learning rates (observational learning; Heyes, 1994) than demonstrators (instrumental learning). Sampaio et al. (2020) and Fiorito & Scotto (1992) observed similar patterns in cuttlefish newborns and common octopus, respectively, supporting the hypothesis that social learning represents an important advantage in preventing energy and time expenditure associated with trial-and-error learning. This concurs in corvids, which exhibit adaptations to sociality through optimized social learning (Templeton et al., 1999).

The increased latency to attack resulting from Severe Hypoxia in demonstrators and observers might signify that cuttlefish will face reduced foraging opportunities during hypoxic events. Indeed, neither activity levels (open-field test) nor reaction times (camouflage latency) were affected by SH, suggesting that hatchlings under SH either display enhanced inhibitory control or, more likely, have reduced motivation to hunt over time (observer cuttlefish spent more time in the experimental arena than demonstrators). This might be due to reduced visual function, which is essential in predatory attacks (Wu et al., 2020). Indeed, hypoxia was found to hinder visual acuity in the fish *Pagrus auratus* (Robinson et al., 2013), but only below critical oxygen tension (P_crit_), at which the animal can no longer maintain oxygen uptake independent from environmental O_2_ levels (Grieshaber et al., 1988). Rosa et al. (2013) reported average critical oxygen tensions in *S. officinalis* embryos of ca. 2.8 kPa (1 mg O_2_ L^-1^), decreasing to an average of 0.15 kPa (0.05 mg O_2_ L^-1^) during the pre-hatching stage. For this reason, we could expect cuttlefish to uphold visual function at hypoxia levels. However, vision oxygen demand grows with the temporal resolution of the eye (Laughlin et al., 1998), which is characteristically high in shallow-water cephalopods (Seibel & Drazen, 2007). Much like the brain, the visual system relies on oxygen-sustained neurons and is further contingent upon photoreceptor cells, which use ATP from oxidative phosphorylation to remain active (Wong-Riley, 2010). Like other marine invertebrates, *S. officinalis* possesses rhabdomeric phototransduction (i.e., uses the photoreceptor rhodopsin instead of rods and cones), which has high ATP demands under lit conditions (Morshedian & Fain, 2017). Additionally, retinal function response is highly species-specific. For instance, retinal impairment in the squid *Doryteuthis opalescens* started at 2 kPa (0.8 mg O_2_ L^-1^), and temporal resolution lessened at 3.8 kPa (1.6 mg O_2_ L^-1^) (McCormick et al., 2019). However, photobehavior is impacted earlier, at 6.4 kPa (2.6 mg O_2_ L^-1^) (McCormick et al., 2022).

A positive control, wherein the cuttlefish was rewarded for its capture attempt at the beginning of demonstrator and observer trials, or activity level assessment at those timestamps, would have been useful to disentangle the causes of decreased attack attempts by SH observer cuttlefish.

### Chronic and acute low oxygen yield similar outcomes on cognitive function

Cuttlefish retained the capacity to learn associatively even under extreme oxygen limitation. Bock et al. (2004) found that acute exposure of 1.5 hours to hypoxia of 2.4 mg O_2_ L^-1^ and 30 minutes of recovery under normoxic conditions does not result in brain tissue deoxygenation in *S. officinalis*. This suggests that cuttlefish are able to protect their brains from hypoxic blood flow stemming from environmental hypoxia if oxygen levels are restored. Yet, any oxygen limitation (chronic or acute) seems to limit learning from observation, as latencies to attack the prey were extended only in observers in normoxic conditions with relation to demonstrators.

In addition, low oxygen increases ambulation, indicating that acute and chronic oxygen loss have similar effects on the cognition of developing cuttlefish if O_2_ levels do not approach hypoxia. In contrast, the goldfish (*Carassius auratus*) decreased speed in response to 1 mg O_2_ L^-1^ hypoxia (Israeli & Kimmel, 1996). The same reduction was observed in Nile tilapia (*Oreochromis niloticus*) at 0.8 mg O_2_ L^-1^, accompanied by a rise in ventilation frequency (Xu et al., 2006).

Although thigmotaxis, i.e., directed movements toward contact, usually with vertical surfaces, was not detected here, Tonkins et al. (2015) showed that cuttlefish exhibit thigmotaxis (referred to as the time spent touching the walls and facing away from them) under stressful conditions (bare plastic or bare glass). Alternatively, cuttlefish display thigmotaxis in the form of ventral adhesion to the arena (von Boletzky & Roeleveld, 2000), which would decrease average speed. Since spatial exploration under hypoxia did not differ from normoxic conditions, conflicting effects of increased activity levels and adhesion to the substrate may have existed. However, it also ensues that thigmotaxis is not responsible for inhibited predatory behavior in hypoxic waters.

Severe hypoxia yielded weaker disruptive patterns. However, the lower pixel intensity range observed was not due to cuttlefish resorting to dark uniform patterns, which is supported by the high minimum intensities observed (denoting an inability to produce dark tones). Interestingly, Court et al. (2022) found that predicted ocean warming and acidification levels jointly elicit an intensification of the disruptive pattern in cuttlefish hatchlings. Consequently, it can be surmised that deoxygenation and hypoxia do not trigger the allocation of energy to defensive behaviors during the cuttlefish’s early ontogeny. Still, it may impair the development of cognitive processes, such as their chromatophore network or visual system, essential for camouflage (Reiter & Laurent, 2020), under extremely low oxygen levels. Visual integrity is paramount, considering that post-embryonic visual experience modulates the maturation of body patterns (*Sepia pharaonis*; Lee et al., 2010). A potential pathway altered by hypoxia is that of glutamatergic transmission through N-methyl-D-aspartate (NMDA) receptor signaling situated in the optic lobe of cuttlefish. It is responsible for experience-dependent behavioral plasticity and, thus, maturation of the visual system and body patterning (Lee et al., 2013). In sand, chromatic patterns were not affected by deoxygenation or hypoxia. This suggests that the mottle pattern is cognitively less demanding than the disruptive pattern.

Hypoxia induces not only direct mortality, but also severe sub-lethal effects such as the inability to hunt and camouflage. Further, we propose that acute and chronic exposure to low oxygen have similar effects on cuttlefish cognition, considering predicted deoxygenation levels and least conservative estimates for hypoxic levels, as demonstrated by the increased activity levels, the ability to learn asocially and inability to learn socially, and the ability to camouflage despite moderate oxygen limitation. As such, irreversible changes are expected to occur from extreme events and interactions between stressors (Cooley et al., 2022). We conclude that expanding hypoxic zones will shrink cuttlefish fundamental niches, as this group will likely be unable to survive in these areas. This study highlights the urgent need for an ambitious cutback on greenhouse gas emissions and control of nutrient discharges into the ocean.

## Funding

British Ecological Society supported this study through a Large Research Grant – LRB21/1004 to JRP. SPECO – Sociedade Portuguesa de Ecologia supported this study through the program “Projetos para Investigadores em Início de Carreira 2020” to JRP. FCT – Fundação para a Ciência e Tecnologia, I.P. supported this study through scientific employment stimulus programs, namely DL57/2016/CP1479/CT0023 to TR and 2021.01030.CEECIND to JRP and under the strategic project UIDB/04292/2020 granted to MARE and project LA/P/0069/2020 granted to the Associate Laboratory ARNET.

## Acknowledgements

We thank Dr. Stefan Mucha for his valuable support during the Ardoxy system set up.

## Notes

### Competing Interest Statement

The authors have declared no competing interest.

