## Supplementary material for "Oxygen loss compromises the survival and cognition of a coastal cephalopod": Table S1

Table S1. Daily SW parameter values (mean ± SD) measured at night (07h30) in each replicate tank during the experiment.

|  | **Normoxia** | **Deoxygenation** | **Mild hypoxia** | **Severe hypoxia** |
| --- | --- | --- | --- | --- |
| **Temperature (ºC)** | 18.05 ± 0.13 | 18.05 ± 0.15 | 18.08 ± 0.08 | 18.06 ± 0.16 |
| **pH** | 8.14 ± 0.11 | 8.09 ± 0.08 | 8.12 ± 0.08 | 8.16 ± 0.09 |
| **Salinity** | 34.95 ± 0.58 | 34.82 ± 0.62 | 34.95 ± 0.63 | 34.96 ± 0.62 |
| **O_2_ (mgL^-1^)** | 7.77 ± 0.11 | 7.02 ± 0.82 | 4.24 ± 0.4 | 2.33 ± 0.85 |
| **O_2_ (%)** | 101.38 ± 1.45 | 91.50 ± 9.23 | 55.31 ± 4.46 | 30.04 ± 6.13 |

Table S2. Daily SW parameter values (mean ± SD) measured during the day (10h00) in each replicate tank during the experiment.

|  | **Control** | **Deoxygenation** | **Mild hypoxia** | **Severe hypoxia** |
| --- | --- | --- | --- | --- |
| **Temperature (ºC)** | 18.13 ± 0.15 | 18,12 ± 0,13 | 18,11 ± 0,11 | 18,1 ± 0,15 |
| **pH** | 8.14 ± 0.1 | 8,05 ± 0,59 | 8,09 ± 0,09 | 8,13 ± 0,1 |
| **Salinity** | 34.91 ± 0.57 | 34,77 ± 0,56 | 34,88 ± 0,62 | 34,92 ± 0,59 |
| **O2 (mg L^-1^)** | 7.98 ± 0.12 | 7,189 ± 0,21 | 7,85 ± 0,13 | 7,78 ± 0,27 |
| **O2 (%)** | 104.18 ± 1.7 | 93,80 ± 2,39 | 102,54 ± 1,77 | 101,64 ± 2,67 |
| ***p*CO_2_ (µatm)** | 463.8 ± 102.4 | | | |
| **TA (µmol/kgSW)** | 2508.6 ± 114.6 | | | |
| **TCO_2_ (µmol /kgSW)** | 2260.86 ± 120.2 | | | |
| **HCO_3_ (µmol /kgSW)** | 2031.52 ± 113.5 | | | |
| **ΩAr** | 2.86 ± 0.71 | | | |

Table S3. Results from type II Wald chi-squared tests based on Cox Proportional Hazards models and Generalized Linear Mixed Models. Effect of treatments (in some cases, number of the trial and interaction, and function – demonstrator or observer) on learning parameters and time-to-hatch. O – Observer, D – Demonstrator.

|  | Predictor | χ2 | d.f. | p-value |
| --- | --- | --- | --- | --- |
| Time-to-hatch | Treatment | 19.3 | 3 | **3e-04** |
| Learning rate | Function | 194.01 | 1 | **< 2.2e-16** |
| Learning success | Function | 22.512 | 1 | **2.089e-06** |
| Classical learning | Treatment | 2.9 | 3 | 0.4 |
| Classical learning (if attack) | Treatment | 2.9 | 3 | 0.4 |
| Social learning | Treatment | 13.17 | 3 | **0.004** |
|  | Function | 22.5 | 1 | **0.000002** |
|  | Treatment (D) | 0.02 | 1 | 0.9 |
| Social learning  (if attack) | Treatment | 5.12 | 3 | 0.2 |
| Latency to first attack | Treatment | 23.0217 | 3 | **3.996e-05** |
|  | Function | 7.9706 | 1 | **0.004754** |
|  | Treatment Trial | 5.2529 | 3 | 0.154187 |

P-values under 0.05 are represented in bold

Table S4. Results from post-hoc multiple comparisons from Cox Proportional Hazards models and Generalized Linear Mixed Models. Time-to-hatch and learning response to the treatments (C – control, DE – deoxygenation, MH – mild hypoxia, SH – severe hypoxia). O – Observer, D – Demonstrator.

| Response |  | Comparisons | df | | | p-value |
| --- | --- | --- | --- | --- | --- | --- |
| Time-to-hatch |  | C - DE | 270 | | | 0.56578 |
|  |  | C - MH | 270 | | | 0.27948 |
|  |  | C - SH | 270 | | | **7.9e-05** |
|  |  | DE - MH | 270 | | | 0.50727 |
|  |  | DE - SH | 270 | | | **0.00392** |
|  |  | MH - SH | 270 | | | **0.00087** |
| Social learning |  | C - DE | 135 | | | 0.6989 |
|  |  | C - MH | 135 | | | 0.5861 |
|  |  | C - SH | 135 | | | **0.0212** |
| Response |  | Comparisons | SE | d.f. | t-ratio | p-value |
| Latency to first attack (D) | | C - DE | 48.6 | 135 | -0.437 | 0.9720 |
|  |  | C - MH | 51.2 | 135 | -1.309 | 0.5591 |
|  |  | C - SH | 57.2 | 135 | -3.510 | **0.0034** |
|  |  | DE - MH | 62.3 | 135 | -0.734 | 0.8831 |
|  |  | DE - SH | 67.3 | 135 | -2.666 | **0.0423** |
|  |  | MH - SH | 69.2 | 135 | -1.932 | 0.2196 |
| Latency to first attack (O) | | C - DE | 55.6 | 135 | 1.418 | 0.4901 |
|  |  | C - MH | 57.2 | 135 | 1.491 | 0.4455 |
|  |  | C - SH | 59.0 | 135 | -1.375 | 0.5169 |
|  |  | DE - MH | 46.7 | 135 | 0.138 | 0.9991 |
|  |  | DE - SH | 48.9 | 135 | -3.271 | **0.0074** |
|  |  | MH - SH | 50.7 | 135 | -3.282 | **0.0071** |
| Latency to first attack (D – O) | | C - C | 50.8 | 267 | -0.501 | **0.0218** |
|  |  | DE - DE | 53.4 | 267 | -1.366 | 0.8717 |
|  |  | MH – MH | 55.9 | 267 | -0.165 | 1.0000 |
|  |  | SH - SH | 65.6 | 267 | -0.480 | 0.9997 |

SE – standard error, d.f. – degrees of freedom, t-ratio – t test statistic. P-values under 0.05 are represented in bold

Table S5. Percentage of cuttlefish that attacked the prawn, that learned (three consecutive trials without attacking), that learned only if they attacked at least once and the sample size (n), in relation to the treatments.

|  | Attacked | Learning | Learning if attacked | n |
| --- | --- | --- | --- | --- |
| Demonstrators | | | | |
| Normoxia | 84.8 | 59.5 | 59.4 | 79 |
| Deoxygenation | 75.0 | 75.0 | 75.0 | 24 |
| Mild hypoxia | 76.2 | 52.4 | 57.1 | 21 |
| Severe hypoxia | 75.0 | 62.5 | 62.5 | 16 |
| Observers | | | | |
| Normoxia | 59.1 | 90.9 | 50.0 | 22 |
| Deoxygenation | 65.9 | 84.1 | 50.0 | 44 |
| Mild hypoxia | 67.6 | 81.1 | 48.6 | 37 |
| Severe hypoxia | 27.3 | 97.0 | 24.2 | 33 |

Table S6. Effect of treatments on open-field test parameters. Results from type II Wald chi-squared tests based on linear models and results from post-hoc comparisons (Tukey-corrected p-values). Responses to the treatments (C – control, D – deoxygenation, MH – mild hypoxia, SH – severe hypoxia).

|  | χ2 | d.f. | p-value |
| --- | --- | --- | --- |
| Average speed | 30.23 | 3 | **1.234e-06** |
| Spatial exploration | 16.915 | 3 | **0.0007359** |
| Wall proximity | 5.5185 | 3 | 0.06063 |

|  | Estimate | | | SE | | d.f. | t-ratio | | p-value | |
| --- | --- | --- | --- | --- | --- | --- | --- | --- | --- | --- |
| Average speed | | | | | | | | | | |
| C - DE | | -31.736 | 0.528 | | 31 | | | -3.286 | | **0.0128** |
| C - MH | | -3.464 | 0.805 | | 31 | | | -4.303 | | **0.0009** |
| C - SH | | 0.277 | 0.574 | | 31 | | | 0.484 | | 0.9622 |
| DE - MH | | -1.728 | 0.815 | | 31 | | | -2.119 | | 0.1693 |
| DE - SH | | 2.014 | 0.594 | | 31 | | | 3.391 | | **0.0098** |
| MH - SH | 3.742 | | | 0.844 | | 31 | 4.433 | | **0.0006** | |
| Spatial exploration | | | | | | | | | | |
| C - DE | | -0.2374 | 0.0752 | | 31 | | | -3.158 | | **0.0176** |
| C - MH | | -0.2903 | 0.0813 | | 31 | | | -3.573 | | **0.0061** |
| C - SH | | -0.0999 | 0.0829 | | 31 | | | -1.205 | | 0.6286 |
| DE - MH | | -0.0529 | 0.0772 | | 31 | | | -0.685 | | 0.9019 |
| DE - SH | | 0.1376 | 0.0799 | | 31 | | | 1.722 | | 0.3300 |
| MH - SH | | 0.1904 | 0.0796 | | 31 | | | 3.392 | | 0.0997 |

P-values under 0.05 are represented in bold

Table S7. Effect of treatments on camouflage responses and measurement validation. Results from type II Wald chi-squared tests based on linear mixed models and results from Pearson correlations between measurements of a photograph subset (20%). Cor – Pearson's correlation coefficient. D.f. - degrees of freedom.

|  | χ2 | d.f. | p-value |
| --- | --- | --- | --- |
| Mantle length | 22.788 | 3 | **7.67e-05** |
| Latency in sand | 9.5753 | 3 | 0.3085 |
| Latency in gravel | 2.6241 | 3 | 0.05338 |
| Burial attempts | 9.3661 | 3 | 0.1043 |
| Sand pixel intensity | 0.0245 | 3 | 0.999 |
| Sand pixel range | 0.6189 | 3 | 0.8921 |
| Gravel pixel intensity | 17.902 | 3 | **0.0004608** |
| Gravel pixel range | 16.914 | 3 | **0.0007361** |
| Measurement validation | | | |
|  | **Cor** | **d.f.** | **p-value** |
| Sand pixel intensity | 0.968 | 18 | **2.793e-12** |
| Sand pixel range | 0.942 | 18 | **5.954e-10** |
| Gravel pixel intensity | 0.978 | 18 | **9.465e-14** |
| Gravel pixel range | 0.973 | 18 | **5.248e-13** |

P-values under 0.05 are represented in bold

Table S8. Results from post-hoc multiple comparisons from regression analyses (linear mixed models, LMM). Camouflage responses to the treatments (C – control, D – deoxygenation, MH – mild hypoxia, SH – severe hypoxia). SE – standard error, d.f. - degrees of freedom.

|  | Estimate | | | SE | | d.f. | t-ratio | | p-value | |
| --- | --- | --- | --- | --- | --- | --- | --- | --- | --- | --- |
| Pixel intensity in gravel | | | | | | | | | | |
| C - DE | | -2.427 | 4.57 | | 85 | | | -0.531 | | 0.9514 |
| C - MH | | -2.753 | 4.64 | | 85 | | | -0.593 | | 0.9339 |
| C - SH | | -16.962 | 4.35 | | 85 | | | -3.901 | | **0.0011** |
| DE - MH | | -0.326 | 4.97 | | 85 | | | -0.066 | | 0.9999 |
| DE - SH | | -14.535 | 4.69 | | 85 | | | -3.098 | | **0.0139** |
| MH - SH | | -14.209 | 4.76 | | 85 | | | -2.986 | | **0.0191** |
| Pixel range in gravel | | | | | | | | | | |
| C - DE | -8.41 | | | 5.38 | | 85 | -1.563 | | 0.4054 | |
| C - MH | 2.51 | | | 5.47 | | 85 | 0.459 | | 0.9676 | |
| C - SH | 13.84 | | | 5.12 | | 85 | 2.703 | | **0.0406** | |
| DE - MH | 10.92 | | | 5.85 | | 85 | 1.868 | | 0.2494 | |
| DE - SH | 22.25 | | | 5.53 | | 85 | 4.027 | | **0.0007** | |
| MH - SH | 11.33 | | | 5.60 | | 85 | 2.021 | | 0.1884 | |
| Mantle length | | | | | | | | | | |
| C - DE | -0.2913 | | | 0.368 | | 84 | -0.792 | | 0.8578 | |
| C - MH | 0.0581 | | | 0.377 | | 84 | 0.154 | | 0.9987 | |
| C - SH | 1.2561 | | | 0.350 | | 84 | 3.589 | | **0.0031** | |
| DE - MH | 0.3494 | | | 0.391 | | 84 | 0.894 | | 0.8082 | |
| DE - SH | 1.5474 | | | 0.370 | | 84 | 4.186 | | **0.0004** | |
| MH - SH | 1.1981 | | | 0.376 | | 84 | 3.187 | | **0.0107** | |

SE – standard error, d.f. – degrees of freedom, t-ratio – t test statistic. P-values under 0.05 are represented in bold

Figure S1. Probability of attacking the prawn-in-a-tube across trials, achieved through time-to-event analysis, under the treatments Normoxia (8 mg O_2_ L^-1^), Deoxygenation (7 mg O_2_ L^-1^), Mild hypoxia (nocturnal 4 mg O_2_ L^-1^) and Severe hypoxia (nocturnal 2 mg O_2_ L^-1^). From demonstrators (D) and observers (O). Lines represent predicted proportions (Cox regression) and shaded areas depict 95% confidence intervals

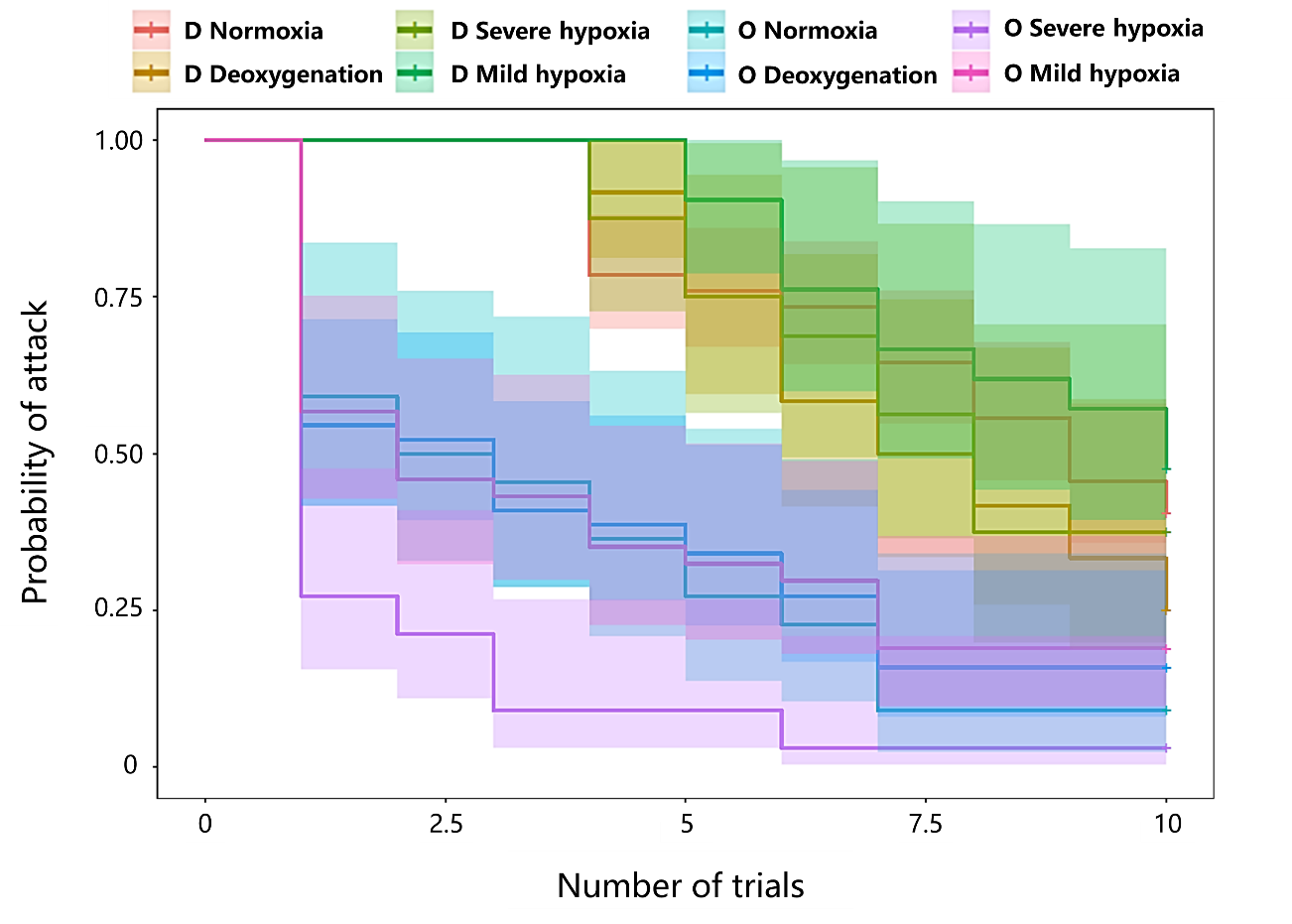

Figure S2. Probability of observers attacking the prawn-in-a-tube across trials, drawn from time-to-event analysis, given the demonstrator is from Normoxia or from their own treatment. Lines represent predicted proportions (Cox regression) and shaded areas depict 95% confidence intervals

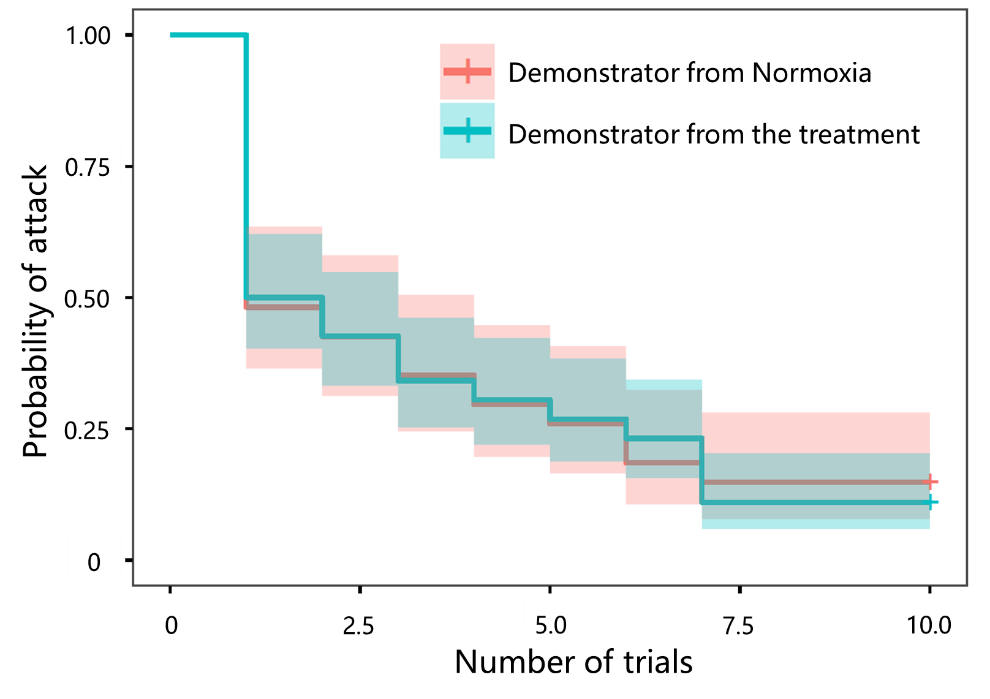

Figure S3. Proportion of space explored during 20 min in the open-field test under the treatments Normoxia (8 mg O_2_ L^-1^), Deoxygenation (7 mg O_2_ L^-1^), Mild hypoxia (nocturnal 4 mg O_2_ L^-1^) and Severe hypoxia (nocturnal 2 mg O_2_ L^-1^). Results are expressed as estimated marginal means ± CI (95%). Comparison significance levels: * p < 0.05, ** p < 0.01, *** p < 0.001

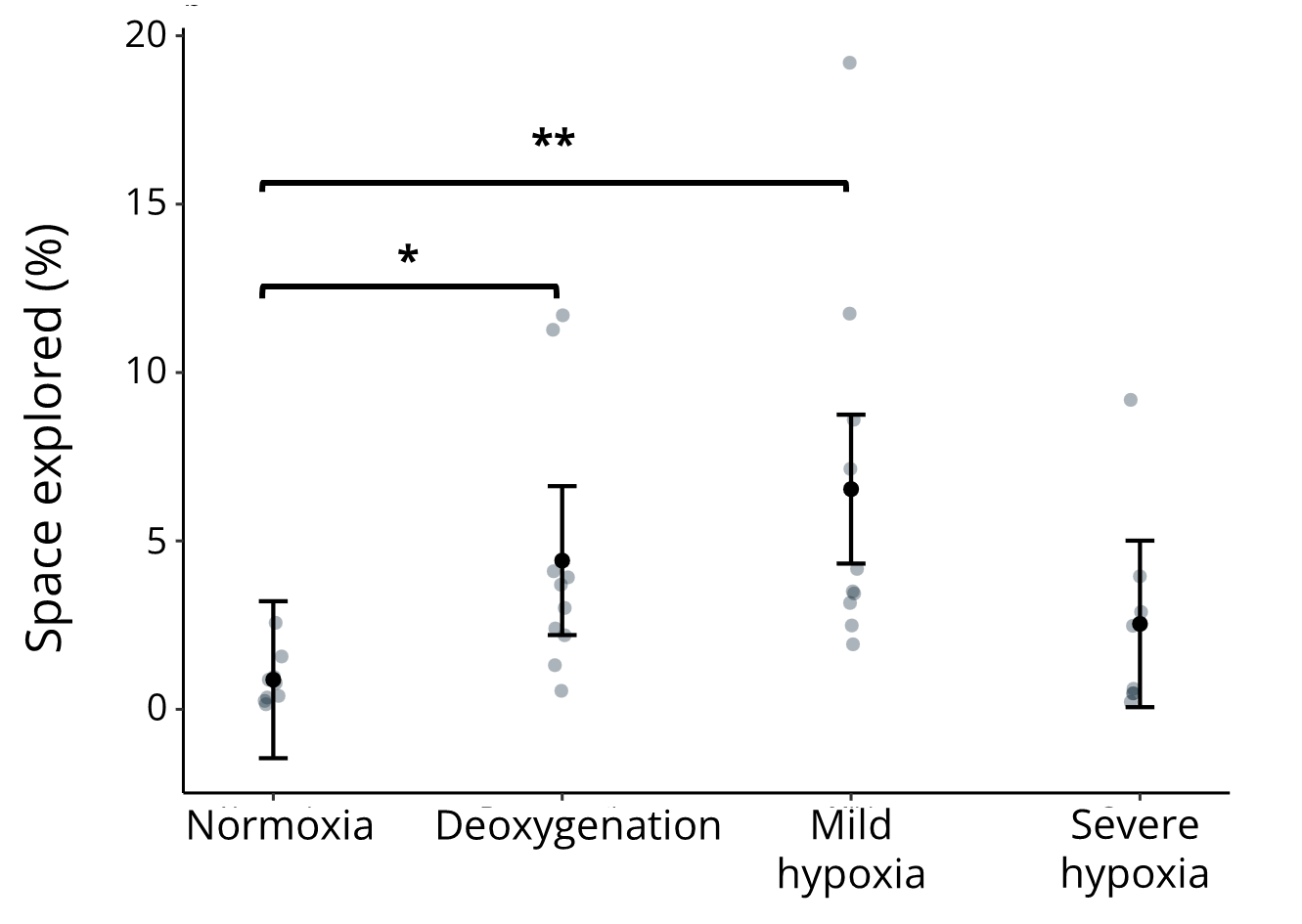

Figure S5. Latency to camouflage under the treatments Normoxia (8 mg O_2_ L^-1^), Deoxygenation (7 mg O_2_ L^-1^), Mild hypoxia (nocturnal 4 mg O_2_ L^-1^) and Severe hypoxia (nocturnal 2 mg O_2_ L^-1^). Results are expressed as estimated marginal means ± CI (95%). Comparison significance levels: * p < 0.05, ** p < 0.01, *** p < 0.001

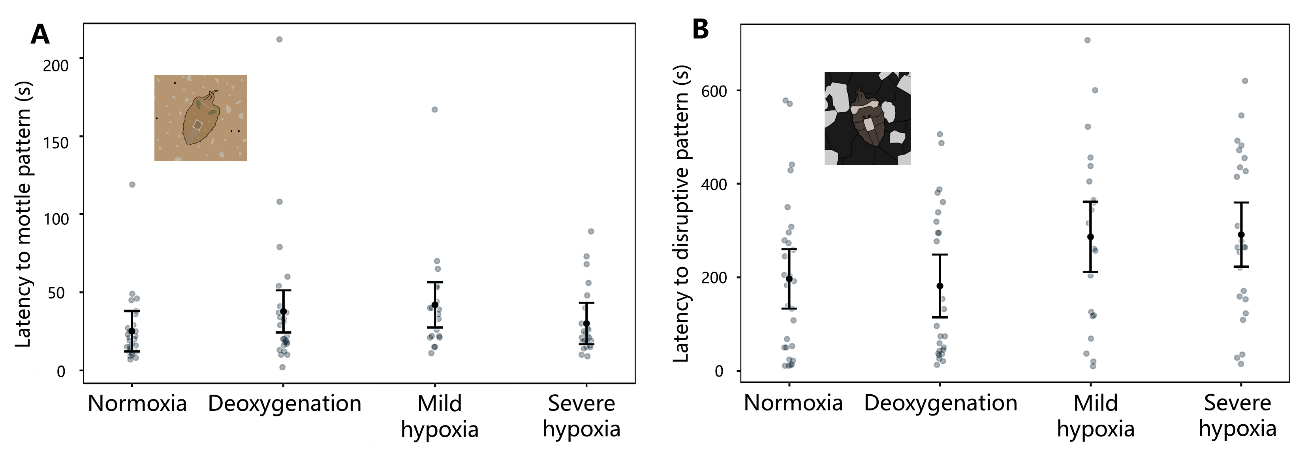
